# A multimodal dataset linking wide-field calcium imaging to behavior changes in mice during an operant lever-pull task

**DOI:** 10.1101/2025.02.03.631599

**Authors:** Masashi Kondo, Keisuke Sehara, Rie Harukuni, Ryo Aoki, Shoya Sugimoto, Yasuhiro R. Tanaka, Masanori Matsuzaki, Ken Nakae

## Abstract

The BraiDyn-BC (Brain Dynamics underlying emergence of Behavioral Change) Database offers an extensive, multimodal dataset that links wide-field calcium imaging of the mouse neocortex to comprehensive behavioral measurements during a behavioral task. As one of the contents in this database, we newly provide a dataset that includes 15 sessions spanning two weeks of motor skill learning, in which 25 mice were trained to pull a lever to obtain water rewards. Simultaneous high-speed videography captures body, facial, and eye movements, and environmental parameters are monitored. The dataset also features resting-state cortical activity and sensory-evoked responses, enhancing its utility for both learning-related and sensory-driven neural dynamics studies. Data are formatted in accordance with the Neurodata Without Borders (NWB) standard, ensuring compatibility with existing analysis tools and adherence to the FAIR principles. This resource enables in-depth investigations into the neural mechanisms underlying behavior and learning. The platform encourages collaborative research, supporting the exploration of rapid within-session learning effects, long-term behavioral adaptations, and neural circuit dynamics.

## Background & Summary

Understanding the relationship between neural activity and behavior remains a fundamental challenge in neuroscience, particularly in the context of learning. Operant conditioning paradigms ^1,2^ have proven invaluable for studying behavioral adaptation. For decades, in combination with conventional neurophysiological approaches, researchers have uncovered fundamental neural correlates of behavioral adaptation ^3–5^. Recent technical advances in wide-field calcium imaging ^6–11^, coinciding with the emergence of deep-learning-based behavioral analysis ^12–15^, offer unprecedented opportunities to correlate cortical activity patterns with precise behavioral quantification across large datasets.

Recent studies have highlighted that even seemingly task-unrelated movements—particularly subtle orofacial ones—can strongly modulate cortical activity of rodents ^16,17^. These covert behaviors and their accompanying neural states can not only shape ongoing behavioral performance, but may also influence longer-term outcomes of learning, underscoring the need for comprehensive datasets capable of capturing both overt and subtle expressions of behavior.

Here we deposit a dataset within BraiDyn-BC Database (https://braidyn-bc-database.netlify.app/), which builds on the earlier studies ^18,19^, addressing the need for comprehensive, multimodal data in learning studies by combining wide-field calcium imaging of cortical activity with detailed behavioral measurements in mice during behavioral tasks under the head-fixed condition. Our experimental paradigm employed transgenic mice expressing GCaMP6s in excitatory neurons ^20^, enabling fluorescent monitoring of neural activity across the dorsal cortex. In addition to behavioral sensors for task control and recording, three high-speed cameras (100 Hz) captured upper body, facial, and eye movements, while the atmospheric environment was continuously monitored. In this newly provided dataset for the database, in which 25 mice completed 15 training sessions of an operant lever-pull task, we successfully collected data from 364 out of 375 sessions, achieving a collection rate of 97%. The 11 sessions could not be measured due to incompleteness of calcium imaging data.

The dataset not only includes task-related measurements but also resting-state cortical activity and sensory-evoked responses under anesthesia, enabling investigation of both task-dependent and baseline neural dynamics. Data is structured in accordance with the Neurodata Without Borders (NWB) format ^21^, ensuring compatibility with existing analysis tools and adherence to the FAIR principles (Findable, Accessible, Interoperable, Reusable) ^22^. By providing ready-to-use preprocessed data in addition to comprehensive raw data (∼17 TB), we facilitate analyses ranging from rapid within-session learning dynamics to long-term behavioral adaptations and associated neural circuit changes. Note that, considering the goal of mapping neural activity to behavior, our approach shares some of the objectives in common with other open, large-scale, cellular-resolution datasets, including those from the International Brain Laboratory (IBL) ^23,24^ and the Allen Brain Observatory (ABO) ^25^.

This resource aims to advance open neuroscience research by enabling investigations into the complex relationships between cortical activity, motor behavior, and environmental factors during learning ^26^. The multimodal nature of our dataset supports diverse analytical approaches, from traditional behavioral metrics to advanced neural circuit analyses. Through open distribution of this comprehensive dataset, we seek to promote transparent, reproducible research and facilitate novel insights into the neural mechanisms underlying behavior change.

## Methods

### Animals

The transgenic mice for wide-field imaging were obtained by crossing VGluT1-Cre mice (B6.Cg-*Slc17a7tm1.1(cre)Hze*/J, Strain #:037512, Jackson laboratory)^27^ and Ai162 mice (B6.Cg-Igs7tm162.1(tetO-GCaMP6s,CAG-tTA2)Hze/J, Strain #:031562, Jackson laboratory) ^28^. All mice were allowed to access food and water *ad libitum* and were housed in a 12:12 hour light-dark cycle (light cycle; 8 AM–8 PM). The mice of both sexes (11 males and 14 females) were used for the experiments. The age ranged from 17–31 weeks at the end of experiments. For at least 3 days before performing the surgery, mice were supplied and habituated with a carprofen-contained sweetened gel (3 g/day, MediGel Hazelnut or DietGel Boost, ClearH_2_O, ME, USA; 0.48 mg/g in gel, Remadile; Zoetis, NJ, USA), which was occasionally provided after the surgery to reduce post-operative pain. All animal experiments were approved by the Institutional Animal Care and Use Committee of the University of Tokyo, Japan.

### Surgical procedures

The surgical procedures were performed according to our previous report (Kondo & Matsuzaki, 2021) with a small modification. Mice were anesthetized by intraperitoneal or intramuscular injection of a ketamine (74 mg/kg; Daiichi Sankyo Propharma, Tokyo, Japan) and xylazine (10 mg/kg; Beyer Pharma Japan, Osaka, Japan) mixture. After anesthesia, an eye ointment (Tarivid; 0.3% w/v ofloxacin, Santen Pharmaceutical, Osaka, Japan) was used to prevent eye-drying and infection. During surgery, body temperature was maintained at 36–37°C with a heating pad. The head of the mouse was sterilized with 70% ethanol and the hair was shaved, and the scalp was incised. After the skull was exposed, the soft tissue on the skull was removed. The temporal muscle was carefully detached from the cranium to improve the observation of temporal cortical areas. A custom head-plate (Misumi, Tokyo, Japan) was attached to the skull using dental resin cement (Estecem II, Tokuyama Dental, Tokyo, Japan). To prevent excitation light from directly entering through gaps between the head-plate and eyes of the mouse, the gaps were filled with a mixture of dental resin cement (Fuji lute BC; GC, Tokyo, Japan) and carbon powder (FUJIFILM Wako Pure Chemical, Osaka, Japan). To prevent drying of the skull surface, a thin layer of cyanoacrylate adhesive (Vetbond; 3M, MN, USA) and dental resin cement (Super bond; Sun Medical, Shiga, Japan) were applied. After the curing of the dental resin layer, UV-curing optical adhesive (NOA81, Norland Products, NJ, USA) was applied several times. This allowed us to observe the cortical activity longitudinally through an intact cranium. The optical adhesive-cured skull was covered with a silicone elastomer (Dent-silicone V, Shohu, Kyoto, Japan) to protect it from dust. An isotonic saline solution with 5% (w/v) glucose and the anti-inflammatory analgesic carprofen (5 mg/kg, Remadile) was injected intraperitoneally after all surgical procedures. The water control schedule was started after at least 3 days of recovery.

### Behavioral apparatus

All behavioral tasks and imaging were performed in darkness inside a dedicated sound-attenuating task box equipped with a lever unit, a licking recorder, a sound presentation system, and a water reward feeder (O’hara & Co., Tokyo, Japan). The atmospheric environment of the task box was monitored during training; temperature, humidity, atmospheric air pressure, and CO_2_ concentration were measured with electronic sensors (BME680, BOSHE Sensortec, Reutlingen, Germany; MH-Z19C, Zhengzhou Winsen Electronics Technology Co., Zehngzhou, China), which were connected to the microcontroller (Seeed XIAO RP2040, Seeed Technology Co., Shenzhen, China) by the I^2^C interface. Outputs of the atmospheric sensors were recorded on the PC at 20 Hz via a USB serial connection. The head-fixed mouse was placed on the body chamber. A lever and an immobile pawrest were set in front of the right and left forelimbs, respectively ^29,30^. The lever was movable but stabilized with a pair of permanent magnets at a base position. A force of 0.04 N was required for the lever-pull initiation ^30,31^. The lever could be pulled up to the distance of 4 mm, where it was physically blocked, and it returns to the base position, due to the magnetic force, without being pulled. The position of the lever was recorded using a rotary encoder (MES-12-2000P, Microtech laboratory, Kanagawa, Japan) equipped at the position 80 mm away from the lever tip. The pulse outputs of the rotary encoder were counted with an NI-DAQ (USB-6229 or PCIe-6321, National Instruments, TX, USA), converted into the arc length, which was recorded with other analog data. A lick spout was placed in front of mice, aligned with their mouths. The licking behavior was monitored by electrically detecting the contact between the tongue and the lick spout, with the signals conveyed to an analog input of the NI-DAQ. All analog voltage inputs, frame synchronization pulses, and the values from the environmental sensor were recorded at 5 kHz sampling rate. Sound cues (10 kHz sinusoidal tone, 70 dB SPL) were presented through a speaker (FT28D, Fostex, Tokyo, Japan) on the left side, 25 cm from the animal. The water reward (4 μL/drop) was fed with a micro pump unit and released from the lick spout mentioned above. The body movement was recorded with a load cell with a rated capacity of 100 g. The displacement detected by the load cell was processed with an instrumental amplifier (LT1167, Analog Devices, MA, USA) into an analog voltage and transmitted to an analog input of the NI-DAQ. Since this analog voltage was not calibrated with an actual weight, the unit was arbitrary. Visual stimuli was emitted from the tip of φ5 mm stainless tube, attached with a red LED on its opposite side, and touch stimuli was generated as a vibration (linear vibration actuator, LD14-002, Nidec, Kyoto, Japan) or an air puff (0.1 MPa) on the whisker pad. The TTL signals controlling reward delivery or sensory stimuli were recorded as an analog voltage with NI-DAQ.

### Behavioral task

Our task schedule spanned approximately one month. First, to facilitate the acclimation of mice to the environment, we conducted 3–8 days of pre-training (see below). On the last pre-training day, we conducted the first resting-state recording session after the pre-training (recording day 0). On the following days, we conducted task training sessions one session per day. There were occasional training breaks of typically 2–3 days without any handling (*e.g.* over weekends; 3.1 ± 0.6 breaks throughout training; 2.3 ± 0.7 days per break; mean ± s.d., *n* = 25 mice). Resting-state recording sessions were also conducted after task training sessions on recording days 1, 7, and 15. Note that, for one animal, the mid-training resting-state session was performed on recording day 8 instead of day 7 (Table 1). One to six days after 15th recording day, the sensory mapping session (recording day 16) was held (Fig. 1c). The entire recording period (recording day 1 to 16) spanned 24.28 ± 3.2 calendar days (mean ± s.d.).

**Figure 1.**
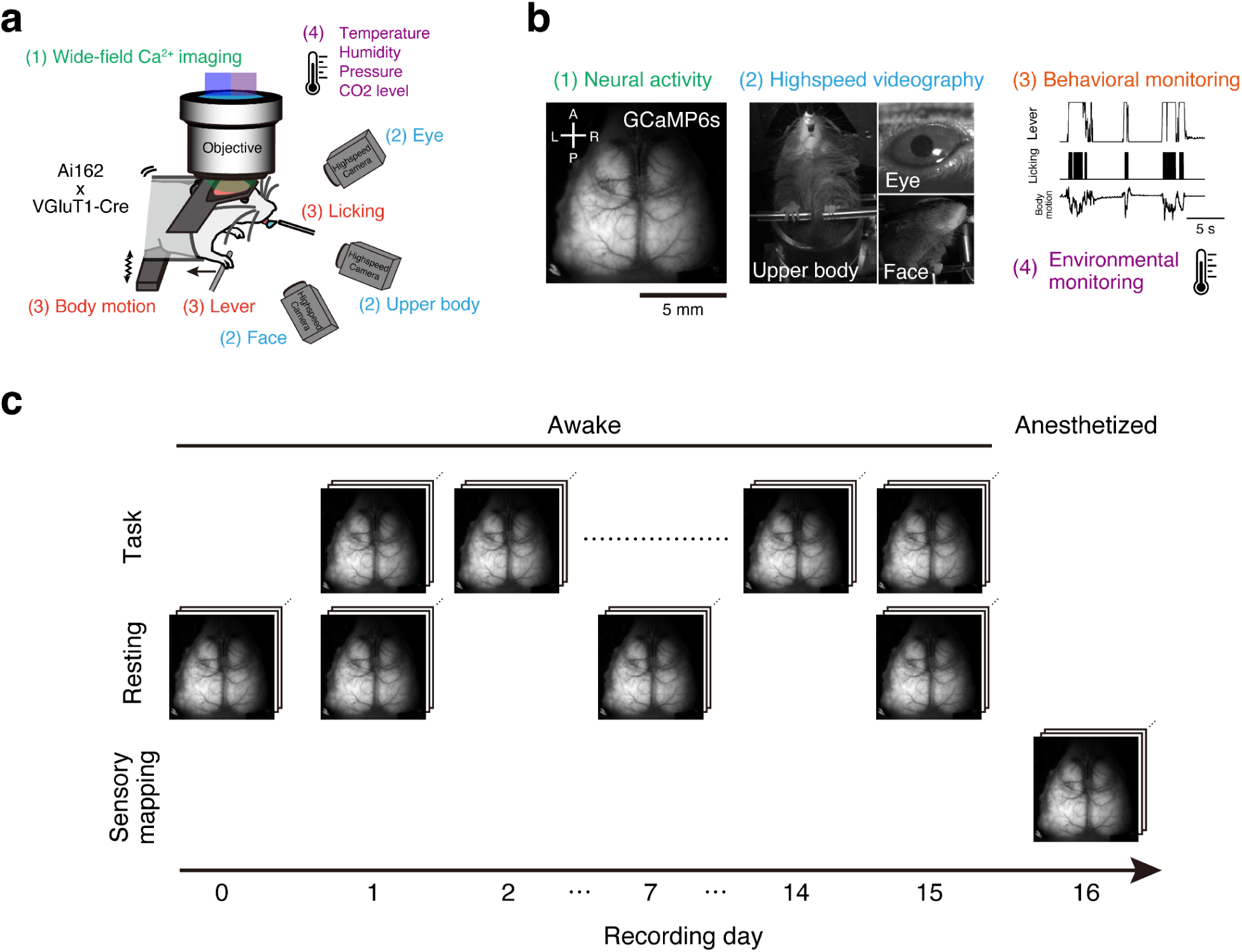
The setup and the timeline of the experiment. **a,b,** Schematic representations of the experimental setup and the data structure. The text colors for setup elements in (**a**) correspond to the colors of the four numbered components in the raw data from typical sessions (**b**). **c**, Timeline of the operant task and the other experiments (See Methods for details).

**Table 1.** Summary of the numbers of different types of sessions reported in this dataset. Out of all the sessions having been performed and recorded, only those accompanying imaging data are deposited here. The denominators on the “Animals” column indicate the total number of animals having been trained for this dataset. The denominators on the “Session” column indicate the total number of sessions having been performed on the corresponding category. The denominators of the other columns correspond to the number of sessions during which imaging data has been successfully recorded. For eight of the animals, two sensory-mapping sessions were performed. Note some sessions experienced failures when recording behavior videos from one or more views. Videos were not recorded during sensory-mapping sessions.

|  | Base recording |  | Sessions with videos |  |  |  |
| --- | --- | --- | --- | --- | --- | --- |
|  | Animals | Session | Upper body | Face | Eye | All views |
| **Task** | 25 / 25 | 364 / 375 | 357 / 364 | 354 / 364 | 356 / 364 | 353 / 364 |
| **Resting state (day 0)** | 19 / 25 | 19 / 19 | 14 / 19 | 14 / 19 | 14 / 19 | 14 / 19 |
| **Resting state (day 1)** | 25 / 25 | 25 / 25 | 25 / 25 | 25 / 25 | 25 / 25 | 25 / 25 |
| **Resting state (day 7)** | 24 / 25 | 24 / 24 | 23 / 24 | 23 / 24 | 23 / 24 | 23 / 24 |
| **Resting state (day 8)** | 1 / 25 | 1 / 1 | 1 / 1 | 1 / 1 | 1 / 1 | 1 / 1 |
| **Resting state (day 15)** | 25 / 25 | 23 / 25 | 23 / 23 | 23 / 23 | 23 / 23 | 23 / 23 |
| **Sensory mapping** | 25 / 25 | 33 / 33 | – | – | – | – |

#### Tone-triggered lever-pull task

The lever-pull task was performed for 30 min per session each day. A lever-pull was defined as the epoch during which the mouse continuously kept the lever over 1 mm pulled from its base position (the maximum pull limit was 4 mm). A trial started with presentation of the sound cue (10 kHz pure tone, 70 dB SPL, 200 ms). When the mouse pulled the lever for longer than a defined duration of time (*T*_pull_) within 1 s from the sound cue onset, these trials were considered successful, and a 4 μL water reward was automatically delivered from the lick spout.

We made sure that the mouse could try pulling the lever only once during the 1 s cued period, by monitoring the lever position on-line. Even before the 1 s had passed for the cued period, once the lever was put into the pulled state, and went back to the non-pulled state before the required duration *T*_pull_ had elapsed, we considered the trial to be a failure, and aborted the trial. On the other hand, if the mouse did not put the lever into the pulled state at all during the cued period, we considered the trial to be a miss.

The inter-trial interval (ITI) between 3–4 s was randomly set after each trial, and if the mice pulled the lever during the ITI, the interval was extended by another 3–4 s until the next sound cue. The required lever-pull time, *T*_pull_, was modulated according to the behavioral performance. The initial *T*_pull_ in the first session was set to 1 ms. It was extended by 50 ms when there was an 80% success rate in the previous 20 trials, up to the maximum duration of 400 ms. We set a refractory period of 20 trials for these extensions, ensuring that each extension occurred at least 20 trials apart. We defined the *T*_pull_final_ of a session as the *T*_pull_ at the end of the session, and the initial *T*_pull_ in the next session was set to *T*_pull_final_ of the previous session minus 100 ms. *T*_pull_ was never shortened within the same session.

#### Resting-state recording

On recording days 1, 7, and 15, resting state recording was conducted after a sufficient intake of water after finishing the lever-pull task. Note that, for one animal, the mid-training resting-state session was performed on recording day 8 instead of day 7 (Table 1). The recording duration was 10 minutes. On day 0, the resting state recording was performed after the pre-training.

#### Sensory mapping

After 15 lever-pull task sessions, mice were allowed to drink water freely in their home cage. After one to six days, a sensory mapping session was performed. The recording time was 15 min under anaesthesia with 0.05 mg/kg fentanyl (Daiichi sankyo company, Tokyo, Japan), 5.0 mg/kg midazolam (Sandoz, Tokyo, Japan) and 0.5 mg/kg medetomidine (Zenoaq, Fukushima, Japan) ^32^. Mice were stimulated visually with a red LED (1 s duration, 0.5 duty cycle, 10 Hz), tactilely with the vibration on the right whisker pad (∼150 Hz; occasionally, with air puff to the whisker pad), and auditorily with the white noise (1 s, ∼70 dB SPL) from the speaker. These stimuli were presented in a fixed order (visual, tactile, auditory), and the inter-stimulus intervals were randomly set to 10–15 s in each trial. After the session, animals were recovered with antagonistic drugs, 0.5 mg/kg flumazenil (Nippon Chemiphar, Tokyo, Japan), 2.5 mg/kg atipamezole (Zenoaq, Fukushima, Japan) and 1.2 mg/kg naloxone (Alfresa Pharma, Osaka, Japan).

#### Water control and pre-training

At least two days before starting training sessions, the water in the home cage was replaced with water containing 2% citric acid^33^. This replacement mildly suppressed the daily water intake of mice. During the period of daily training, mice typically obtained the necessary water (>1 mL) in training sessions, and they had unrestricted access to standard pellets in their home cages. When a training break lasted two or more days (*e.g.*, over weekends), water containing 2% citric acid was placed in the home cage. These bottles were removed the morning of the next training day.

For acclimating to the head-fixed condition in the task box, mice were pre-trained. In pre-training, mice were head-fixed and received water rewards in a way similar to the lever-pull task, except they obtained the reward by licking the lick spout rather than pulling the lever. Pre-training was conducted 3–8 times, and the number of pre-training was included in the metadata. We noticed that some mice gradually lost weight over days even if the mice received daily water intake in the behavioral session. Therefore, to maintain the body weights of mice at approximately > 80% of body weight before the water restriction, we provided mice with additional water and high caloric food (Calorie-Mate fruit flavor, Otsuka Pharmaceutical Co., Tokyo, Japan) when necessary after the behavioral session.

### Acquisition of wide-field one-photon images

For wide-field one-photon calcium imaging, a wide-field tandem-lens macroscope (THT mesoscope, Brain vision, Tokyo, Japan), equipped with the objective lens (PLAN APO 1×, #10450028, Leica microsystems, Wetzlar, Germany) and the imaging lens (F2.0, focal length 135 mm, Samyang, Seoul, Republic of Korea), were used. Images were acquired with a CMOS camera (ORCA FusionBT, C14440-20UP, Hamamatsu photonics, Shizuoka, Japan) by the control of HCImage software (Hamamatsu photonics). Single images consisting of 588 × 588 pixels were captured at 60 Hz. We alternately illuminated two excitation LEDs with different wavelengths (405 nm and 470 nm; M405LP1 and M470L5, Thorlabs, NJ, USA). The lights from the LED sources were cleaned with bandpass emission filters (FBH405-10 for 405 nm and FBH470-10 for 470 nm, Thorlabs). These lights were combined with a dichroic mirror (DMLP425R, Thorlabs) and delivered to the macroscope through a liquid light guide (Ø5 mm Core) and a collimator (LLG5-6H and COP1-A, Thorlabs). The collimated light was passed through the 3D-printed field stop (the geometry was designed for the inner space of the head-plate) and the condenser lens (plano convex lens, f=150 mm, LA1417, Thorlabs), a dichroic beam splitter (FF484-FDi01, Semrock; IDEX Health & Science, NY, USA), then projected to the sample specimen. When performing calcium imaging, a 3D-printed light shield was placed on the head-plate to prevent direct illumination of excitation light to the eyes of the mouse. The total power of excitation lights (blue and violet) was set at ∼10 mW and the fluorescent signal degradation across imaging sessions was not observed. Fluorescent emission signal from the sample was collected with the objective and passed through the dichroic, bandpass filter (FF01-536/40, Semrock; IDEX Health & Science), imaging lens, then projected to the CMOS camera. Since two sequential images obtained by two different excitation lights are analyzed as a pair (see below), the effective sampling rate should be considered 30 Hz. LED illumination and image acquisition timings were recorded using the same DAQ device (USB-6229 or PCIe-6321) used for the task. We imaged 108000 frames (30 min) in each task session, 36000 frames in each resting-state session (10 min on recording days 1, 7, 15), and 54000 frames in each sensory mapping session (recording day 16).

### Acquisition of high-speed videography

Three machine vision cameras (acA1440-220um, Basler, Ahrensburg, Germany) took videos of the upper body, the right side of the face, and the right eye at 100 Hz. Pulses (5V, 100Hz, duty rate of 0.5) from the DAQ device were used for synchronization of the frame acquisition across the cameras, and were recorded as an analog voltage. The bandpass IR cut filter (center 850 nm, 25 nm FWHM, Edmund optics, NJ, USA) was placed in front of an imaging lens of each camera. The videos were saved as MP4 files. LED arrays (OSI3CA5111A, OptoSupply, Hong Kong; consisting of 850 nm 64 LEDs) at the back of each camera were used as light sources. In addition, a 910 nm LED was lit in front of the mouse. The brightness of the 910 nm LED was modulated session-wise, so that the pupil diameter would dilate moderately during imaging, without any adjustments during individual sessions.

### Data processing

Behavioral data processing (filtering, event detection, lick-rate calculation, temporal down-sampling to synchronize for calcium imaging data; see next section) and imaging data processing (spatial down-sampling, interleaved image separation of two excitation wavelength, motion correction; see next next section) were conducted with MATLAB software suite (2022b; Mathworks, MA, USA). All other data processing was conducted in a Python environment with appropriate software packages. The source codes of all processing workflows are deposited at: https://github.com/BraiDyn-BC/bdbc-data-pipeline.

#### Event detection from behavioral sensors and the synchronization for calcium imaging data

Each task event (onsets of tone-cue, reward, lever-pulling, and sensory-stimuli in the sensory mapping experiment) was detected by appropriate threshold. The motion sensor output was processed by applying a 10 Hz cutoff low-pass filter and then subtracting its time-average. A 0-1 function indicating the voltage rise time points in the licking sensor was convolved with a 0.5 s exponential kernel to yield the lick rate. Raw data sampled at 5 kHz were embedded into the 30 Hz imaging frames as follows: any sound cue, reward delivery, or sensory stimulus occurring between the end of one frame and the beginning of the next was assigned to the earlier frame during down-sampling. Lever position, lick rate, and environmental sensor values were averaged over the interval from the onset of a frame to the onset of the following frame to generate the values for the preceding frame.

#### Image processing, registration to standard brain atlas, hemodynamics correction, and calcium activity extraction from wide-field one-photon imaging data

Images were down-sampled into 288 × 288 pixels and divided into two image stacks according to the recorded LED pulse timings. One divided image stack contained images acquired with blue light excitation (*I*_B_) and another with violet light excitation (*I*_V_). The displacement of each frame of *I*_B_ was estimated with the NoRMCorre-based rigid frame registration ^34^ using the time-averaged image of *I*_B_ as a reference. The calculated displacement vector was applied to both *I*_B_ and *I*_V_ stacks, and these motion-corrected *I*_B_ and *I*_V_ stacks are treated as the raw data in the datasets.

To draw neocortical areal borders on our imaging data, we first prepared a “template frame” for each animal, and aligned Allen common coordinate framework (Allen CCF) ^35^ with this template frame using an approach based on MesoNet ^36^. The animal-by-animal template frames were computed by alignment of the session-average frames with each other; for each session, we first computed the mean of the 470 nm-excitation frames over time. We then selected one of these session-mean images as the animal-representative image, and estimated affine transformation matrices for conversion to this representative image from the other session-mean images, based on the keypoints detected using the Oriented FAST and rotated BRIEF (ORB) descriptor ^37^. Using these affine matrices, individual session standard-deviation images over time were warped, and then averaged to obtain the animal-by-animal template frames. The landmark-inference DeepLabCut ^12^ network from MesoNet was used to estimate the nine landmarks on the skull of the animal-by-animal template frames. To estimate the affine transformation from Allen CCF to each template frame, the landmarks being estimated with the likelihood above 0.85 were aligned with those defined on the Allen CCF (provided by the authors of MesoNet). Finally, based on the two affine matrices (i.e. the atlas to the animal template, and the animal template to the session mean), we computed the transformation from Allen CCF to each session-mean image. The ROI (region of interest) masks, being provided with the MesoNet package, were transformed using this final affine matrix, to generate corresponding binary masks of neocortical areas. The signal intensity was averaged across ROI into a time series of signals.

The signals obtained from ROIs were corrected for hemodynamics effects in the similar manner as the others ^17^. Fluorescent signals obtained with blue light (*F*_B_) contained mainly calcium-dependent signal changes but with small contamination of fluctuations due to blood flow changes. Fluorescence obtained with violet light (*F*_V_) could be seen as an instant reference of blood flow changes because its excitation wavelength was near the isosbestic point of GCaMP. We thus correct *F*_b_ based on *F*_v_ by using the following procedure. First, ratiometric signals were calculated as Δ*F*_B_/*F*_B_ = (*F*_B_ - *F*_B0_) / *F*_B0_ and Δ*F*_V_/*F*_V_ = (*F*_V_ - *F*_V0_) / *F*_V0_, with *F*_B0_ and *F*_V0_ being the median of *F*_B_ and *F*_V_ over time, respectively. The two time series, Δ*F*_B_/*F*_B_ and Δ*F*_V_/*F*_V_ were then band pass-filtered to obtain (Δ*F*_B_/F_B_)_filtered_ and (Δ*F*_V_/*F*_V_)_filtered_, to remove high-frequency acquisition noise and low-frequency baseline drifts. In doing so, the “filtfilt” function of SciPy (version 1.14.0) was applied in combination with the fifth-order 0.01–10 Hz band-pass butterworth filter. Finally, linear regression was performed to predict (Δ*F*_B_/F_B_)_filtered_ based on (Δ*F*_V_/*F*_V_)_filtered_, to estimate the hemodynamics effects (Δ*F*_B_/F_B_)_hemo_ = *A* × (Δ*F*_V_/*F*_V_)_filtered_ + *b*, with *A* being the slope and *b* being the bias term. The hemodynamics-corrected calcium signal was computed as the residuals of the regression, i.e. (Δ*F*_B_/F_B_)_corrected_ = (Δ*F*_B_/F_B_)_filtered_ - (Δ*F*_B_/F_B_)_hemo_.

#### Motion estimation of body-parts from high-speed videography

We used DeepLabCut 2.3.10 ^12^ to estimate keypoints representing body-part positions on the behavioral videos (Fig. 2; Table 2). The strategy for extraction of images from videos for neural-network model preparation is described as Fig. S3, with the number of images used being summarized in Table 3 (also see Fig. 5). To improve generalization of some models, we additionally used some videos from different experiments that use the same behavioral rig for initial training iterations. For later training iterations, training and testing of the models were performed in an incremental manner; we verified the performance of the model in tracking a set of videos, before extracting frames from another set of videos. Frames were not extracted from the videos if we considered the tracking performance of the model to be satisfactory. Prior to each training iteration, an image augmentation step was inserted, by using the “imgaug” Python package, to produce 10–20 augmented images based on each of the annotated images to be used for training and testing. The coordinates were set so that the x-axis increases as points move to the right, and the y-axis increases as points move downward. Keypoint estimation by DeepLabCut was inherently performed even when the body part in question was absent. Therefore, in our published data, we shared not only the raw estimations from DeepLabCut but also the likelihood of the estimation for the potential filtering purposes. To estimate the size and the position of the pupil, the set of circumferential points along the pupil boundary was fitted with an ellipse on a frame-by-frame basis. The center position and the length of the major axis of the fitted ellipse were defined as the position and the diameter of the pupil, respectively.

**Figure 2.**
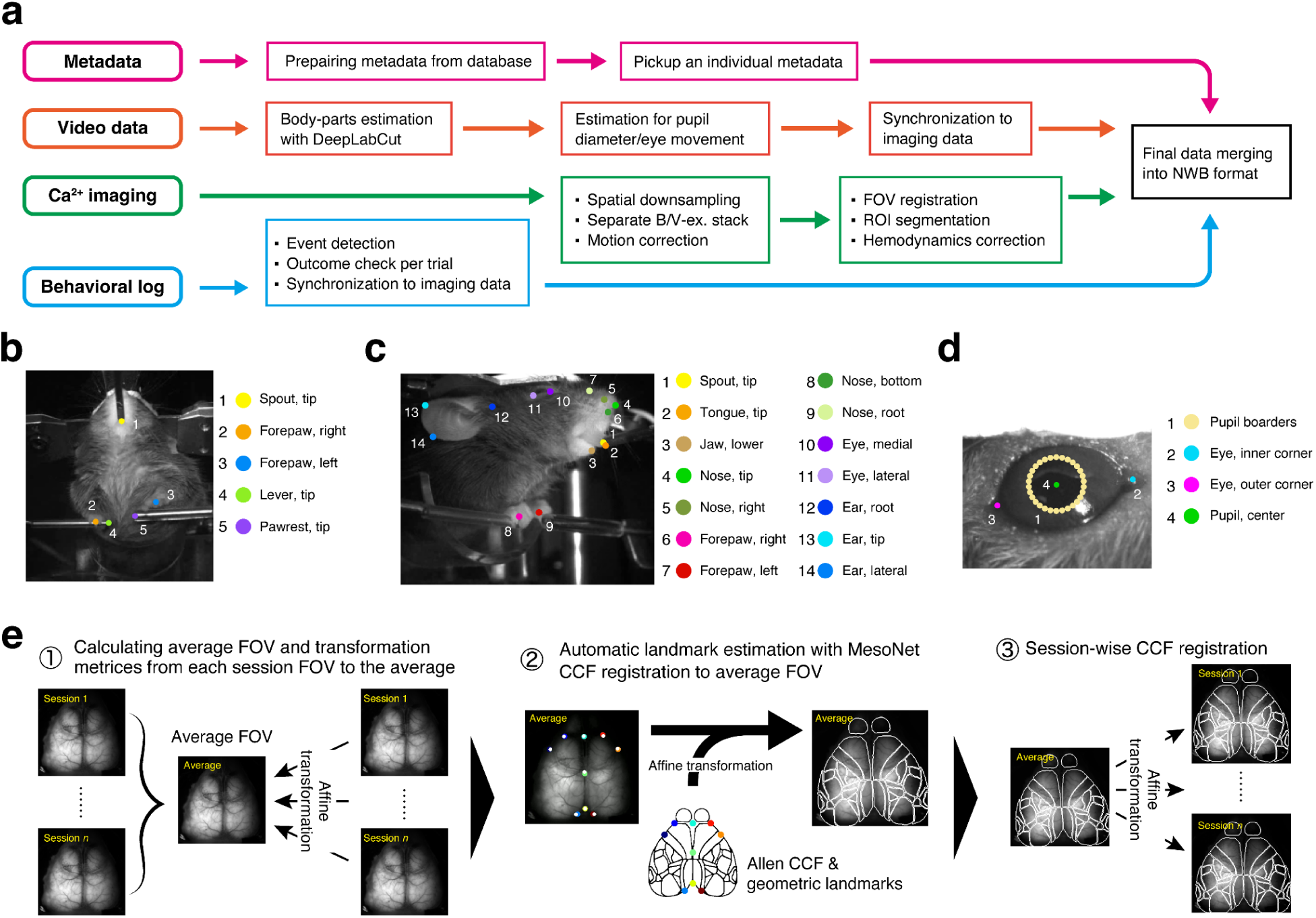
The analytical pipeline of the dataset. **a**, Schematic illustrations of the analytical pipeline. FOV is short for field of view. **b–d**, Keypoints on videos of the upper body (**b**), videos of the right side of the face (**c**), and videos of the right eye (**d**) detected by DeepLabCut. These keypoints are also summarized in Table 2. **e**, Schematic illustration of atlas registration to Allen CCF using MesoNet and affine transformations (see Methods).

**Table 2.** Summary of keypoints (Fig. 2b–d). *This keypoint was not directly estimated by DeepLabCut, but through fitting of an ellipse to the pupil edges (see Methods).

| Keypoints on videos of the upper body |  |  |
| --- | --- | --- |
| 1 | Spout, tip | The tip of the lick spout. |
| 2 | Forepaw, right | The center of the right forepaw. |
| 3 | Forepaw, left | The center of the left forepaw. |
| 4 | Lever, tip | The tip of the lever |
| 5 | Pawrest, tip | The tip of the pawrest |
| Keypoints on videos of the right side of the face |  |  |
| 1 | Spout, tip | The tip of the lick spout. |
| 2 | Tongue, tip | The tip of the tongue in the midline. |
| 3 | Jaw, lower | The tip of the lower jaw in the midline. |
| 4 | Nose, tip | The nose tip in the midline. |
| 5 | Nose, right | The most lateral point of the right nostril wall. |
| 6 | Forepaw, right | The center of the right forepaw. |
| 7 | Forepaw, left | The center of the left forepaw. |
| 8 | Nose, bottom | The most ventral point of the nose in the midline. |
| 9 | Nose, root | The nose root in the midline. |
| 10 | Eye, medial | The inner corner of the right eye. |
| 11 | Eye, lateral | The outer corner of the right eye. |
| 12 | Ear, root | The most anterodorsal tip of the right ear root. |
| 13 | Ear, tip | The most lateral point of the right ear helix. |
| 14 | Ear, lateral | The ventrolateral midpoint of the right ear helix. |
| Keypoints on videos of the right eye |  |  |
| 1 | Pupil borders | Thirty circumferential points on the pupil |
| 2 | Eye, inner corner | The medial corner of the right eye. |
| 3 | Eye, outer corner | The lateral corner of the right eye. |
| 4 | Pupil, center* | The center of the right pupil. |

**Table 3.** Summary of training and validation of the DeepLabCut models. Training of the models was performed incrementally, through checking video files one-by-one. The left-hand side of the numbers of animals, sessions and images for training and testing corresponds to the numbers from the videos of the current dataset, whereas the right-hand side indicates the numbers from different experiments in the same behavioral rig, with the same video-camera configurations. For more details of our testing and validation strategy, see Fig. S3. For the eye model, in particular, discrepancy between the numbers of extracted and annotated frames occurs due to extraction of images when the animals were blinking.

|  | Training and testing |  |  |  | Validation |  |  |
| --- | --- | --- | --- | --- | --- | --- | --- |
|  | *Animals* | Extracted<br>*Sessions* | *Images* | Annotated | *Animals* | Annotated<br>*Sessions* | *Images* |
| **Upper body** | 19 + 4 | 35 + 8 | 1360 + 40 | 1399 | 25 | 96 | 192 |
| **Face** | 6 + 0 | 12 + 0 | 240 + 0 | 240 | 25 | 96 | 192 |
| **Eye** | 23 + 4 | 39 + 8 | 720 + 40 | 676 | 25 | 91 | 180 |

For the resampling of the positional time series from the videography frame rate (100 Hz) to the effective imaging frame rate (30 Hz), we first up-sampled the series to the NI-DAQ sampling rate (5 kHz) and then down-sampled them to 30 Hz. During up-sampling, we dropped the inferred keypoint positions with their likelihood being below 0.2, and plugged the remaining values into the durations of their corresponding video pulses in the 5 kHz NI-DAQ recording.

Inter-pulse interpolation was then performed only when the two neighboring pulses contained valid values. As the imaging pulses, neighboring two LED pulses (one violet and one blue) were merged to obtain the 30 Hz pulses. The mean over the duration of each of these pulses was computed to obtain the series down-sampled to the imaging frame rate.

## Data Records

The dataset files in NWB format are accessible at the repository https://drive.google.com/drive/folders/1yiu-4C3_54SENCy-OLp-E9VgGUgeh8Hd?usp=drive_li <u>nk</u>. The complete dataset with all the raw-data files will be uploaded in the repository https://mmlab-repo.m.u-tokyo.ac.jp/r/Matsuzaki/BraiDyn-BC_CuedLeverPull/. These repositories will be made public with DOIs after the acceptance of this paper.

Our dataset includes data from the 15 behavioral task sessions and the 4 resting-state recording sessions. Each experimental condition and session provides data in both raw and processed forms (see below). For each session, the dataset comprises one NWB file, two TIFF files, and three MP4 files. This data packaging into the NWB format was conducted with a Python environment.

### NWB Files

The NWB file contains both raw and processed data, as well as associated metadata as shown in Table 4. Processed data include:

1. Imaging: calcium activity data segmented into anatomical regions defined by the Allen CCF (22 ROIs per hemisphere).
2. Estimated positions of body parts/pupil: Body-part and pupil coordinates extracted by DeepLabCut (DLC) from the three video recordings. We provide the data at the original video frame rate, as well as those down-sampled to the imaging frame rate.
3. Behavioral data: The data having been synchronized and processed according to the imaging frame timestamps. The NWB file also contains the corresponding raw behavioral data channels.

**Table 4.** Summary of the behavioral and sensor data being stored in each NWB file along with the imaging and video data. DS, downsampled data. *For these signal types, binary values, indicating the occurrence and duration of individual events, are stored instead of the original TTL signals. ^#^For reward-delivery pulses, binary values, indicating the onsets of individual reward-delivery events, are stored instead of the original TTL signals.

| Name | Raw | DS | Unit | Type | Description |
| --- | --- | --- | --- | --- | --- |
| tone | ✓ | ✓<sup>(*)</sup> | V | Digital (TTL) | The pulses representing when the auditory cue (10 kHz pure tone) is on. |
| lever | ✓ | ✓ | mm | Analog | The distance of the lever from its baseline position. |
| reward | ✓ | ✓<sup>(#)</sup> | V | Digital (TTL) | The pulses used for reward delivery. |
| lick | ✓ | ✓<sup>(*)</sup> | V | Digital (TTL) | The output of the lick sensor indicating when the animal licked the spout. |
| lick_rate | – | ✓ | Hz | Analog | The lick rate calculated by applying an exponential kernel to the lick-event onset series. |
| motion | ✓ | ✓ | a.u. | Analog | The output of the load-cell motion sensor. |
| pull_duration | ✓ | – | ms | Integer | The minimum duration for the animal to keep pulling the lever during the trial at the given time point. |
| state_lever | ✓ | ✓ | NA | Boolean | The binary status indicating whether the lever is in the pulled state. |
| state_task | ✓ | ✓ | NA | Integer | The state of the task: waiting phase (0), cued phase (1) or reward phase (2). |
| air_pressure | ✓ | ✓ | hPa | Analog | The atmospheric air pressure level around the behavioral setup. |
| CO2_level | ✓ | ✓ | ppm | Analog | The atmospheric carbon dioxide level around the behavioral setup. |
| humidity | ✓ | ✓ | % | Analog | The atmospheric humidity around the behavioral setup. |
| room_temp | ✓ | ✓ | °C | Analog | The atmospheric temperature around the behavioral setup. |
| LED_B | ✓ | – | V | Digital (TTL) | The pulses representing when the blue (470 nm) LED is on. |
| LED_V | ✓ | – | V | Digital (TTL) | The pulses representing when the violet (405 nm) LED is on. |
| img_acquisition | ✓ | – | V | Digital (CMOS) | Pulses from the camera for calcium imaging, each of which corresponds to the timing when a single frame becomes ready. |
| video_trig | ✓ | – | V | Digital (TTL) | The pulses representing when video-frame acquisition should be started. |

The NWB file includes the `session_description` identifier, `timestamps_reference_time` (timestamp reference, or session start time), and general metadata (`session_id`, `experimenter`, `lab`, `institution`, as well as subject information such as `age`, `genotype`, `sex`, `species`, `subject_id`, `weight`, `date_of_birth`, `strain`), and device information. All data channels are stored under `acquisition` (for raw data) and within `analysis` or `processing` (for processed data). The `analysis` group includes `atlas_to_data_transform`, an affine transformation matrix mapping the Allen CCF atlas (512 × 512 pixels) to the acquired imaging data (288 × 288 pixels).

In processing/behavior/data_interfaces, DLC-extracted keypoints are stored (eye_video_keypoints, face_video_keypoints, body_video_keypoints), along with eye_position (center_x and center_y) and pupil_tracking. The downsampled/data_interfaces/trials group contains trial information synchronized with imaging, as well as the downsampled sensor data (CO2_level, air_pressure, humidity, lever, lick lick_rate, motion, reward, room_temp, state_lever, state_task, tone).

### Imaging Raw Data (TIFF Files)

1. Blue-excitation image: `(animal)_(date)_(task/resting-state)-day(day#)_B.tif`
2. Violet-excitation image: `(animal)_(date)_(task/resting-state)-day(day#)_V.tif`

These TIFF files contain wide-field one-photon imaging data of dorsal cortical calcium activity in mice expressing Ca^2+^ probes in excitatory cortical neurons. During recording, blue and violet excitation lights were alternately applied at 60 Hz to measure both calcium-dependent and calcium-independent fluorescence signals. After data acquisition, frames were separated by excitation wavelength, motion-corrected, and spatially downsampled. The resulting data are stored as 288×288-pixel TIFF files at (recording duration in seconds × 30 Hz) frames.

### Videography Raw Data (MP4 Files)

1. Body Camera Movie: `(animal)_(date)_(task/resting-state)-day(day#)_body.mp4`
2. Face Camera Movie: `(animal)_(date)_(task/resting-state)-day(day#)_face.mp4`
3. Eye Camera Movie: `(animal)_(date)_(task/resting-state)-day(day#)_eye.mp4`

These MP4 files capture the mouse’s upper body, right facial region, and right eye at 100 Hz using three cameras. The cameras are synchronized based on a 5 V, 100 Hz pulse train (50% duty cycle) from a D/A converter. This pulse train is recorded as analog data and can be used to synchronize the videos with both behavioral and calcium activity data.

### Technical Validation

We provided the dataset containing 1) cortical calcium activity obtained with one-photon wide-field imaging, 2) video-tracked body-part movements, 3) sensor-monitored behavioral measurements, and 4) atmospheric environmental parameters during an operant lever-pull task (Fig. 1a,b; Table 1). The dataset incorporates not only 15 task training sessions but also four resting-state recording sessions and one sensory mapping session per animal (Fig. 1c). The data were processed and integrated into session-wise NWB files by the newly developed analysis pipeline, which is available in a public Github repository (Fig. 2). We included in the dataset both raw and processed data, aiming to facilitate for users to utilize it quickly for their purpose and, at the same time, to ensure that they can analyze the data from scratch and go even beyond our scope. We validated the structure of each session-wise data by a test code, which is also provided in the Github repository.

During an operant lever-pull task (Fig. 3a,b), mice (14 female, 11 male) showed various behavioral changes. The response rate to the sound cue tended to increase overall (∼0.55 in task training session 1 to ∼0.85 in task training session 15; median), while the success rate seemed to be constant as a whole (Fig. 3c). To earn rewards efficiently, developing the skill to pull the lever for a long duration (up to 400 ms) while maintaining a high success rate is a straightforward strategy. Therefore, we analyzed *T*_pull_final_ and found that two distinct trends of change in *T*_pull_final_ over the course of task training sessions (Fig. 3c,d). The hierarchical clustering revealed that mice could be divided into two clusters in terms of *T*_pull_final_ (Fig. 3d). Mice belonging to one cluster (cluster A) showed high success rates in the early sessions (∼5) leading to high *T*_pull_final_ which were often maintained even in the late sessions. Mice belonging to the other cluster (cluster B) rarely achieved a high *T*_pull_final_, while they accomplished a response rate and success rate similar to the cluster A mice in the late sessions (Fig. 3e, Fig. S1). These visualizations validate that the present dataset captures various behavior changes during task training sessions. Of the 375 task training sessions conducted, we successfully collected data from 364 sessions (Fig. 3d; Table 1) and completed data collection for all 15 sessions in 16 out of 25 animals.

**Figure 3.**
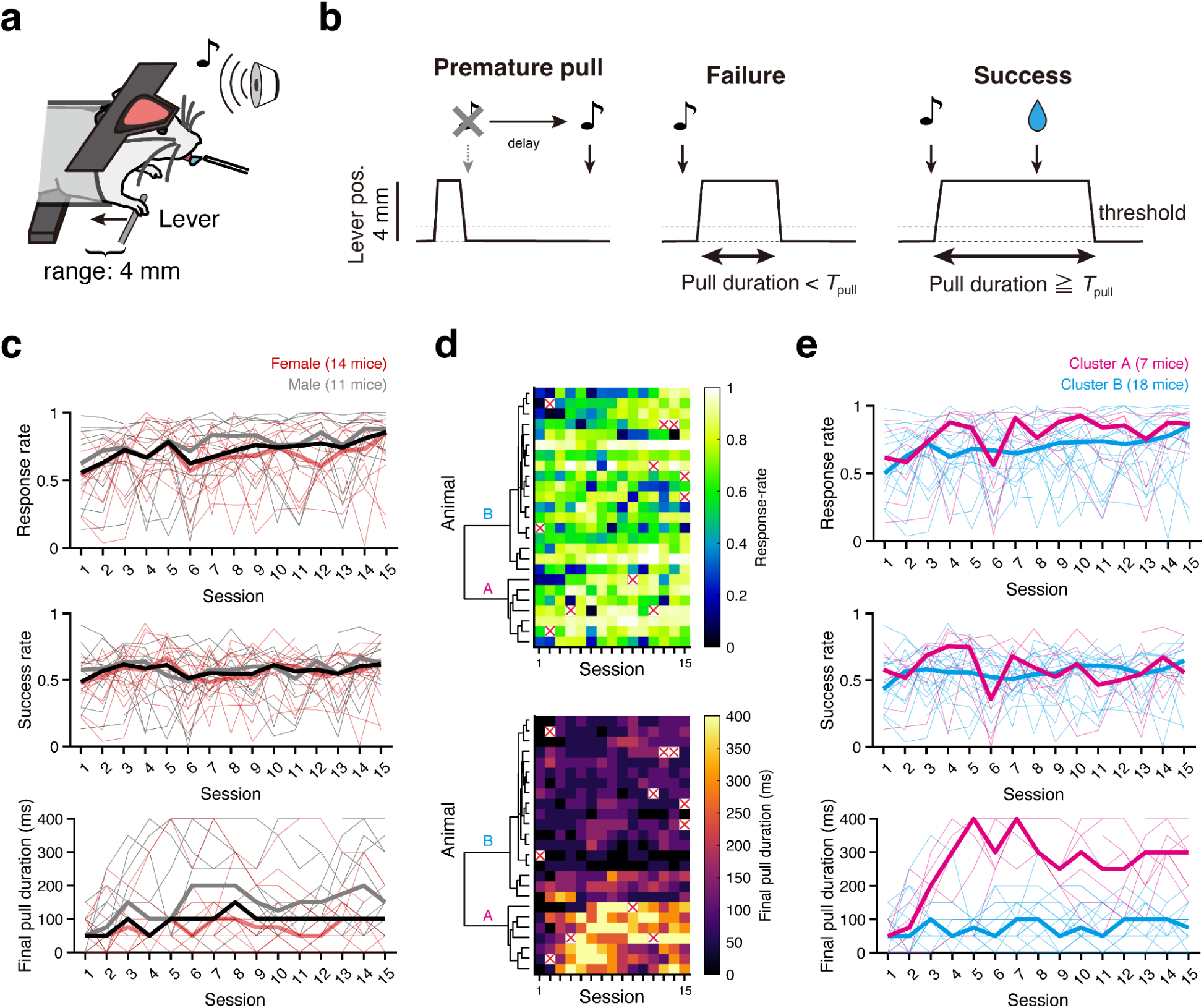
Behavioral performance in task training sessions. **a,b,** Schematic illustrations of the lever-pull task for head-fixed mice are shown. In response to the sound cue, mice should pull the lever and hold it for a specified duration to receive a water reward from the lick spout (**a**). Premature pulls before the sound cue caused an additional delay in sound cue presentation (left panel). Lever pulls initiated within 1 s after the sound cue were considered valid, and a valid pull that exceeded 1 mm (threshold) for the predetermined duration (*T*_pull_; See Methods for details) was considered as successful and rewarded (right panel). An insufficient pull (middle panel) or no response within 1 s after the sound cue was considered a failure. **c,** Response rate (the number of valid pulls/the number of the sound cue; top panel), success rate (the number of successful pulls/the number of the sound cue; middle panel), and final pull duration (*T*_pull_final_; see Methods for details; bottom panel) across sessions are shown. Thin lines show performances of individual animals (red, female; gray, male), whereas thick lines are the median across the population (red, female; gray, male; black, all). **d**. Color-coded response rate and *T*_pull_final_ across sessions are shown. In some sessions, data were missing for various reasons (red crosses, the overall ratio: 11/375 = 0.29%). The result of hierarchical clustering (Ward’s method) based on the chronological change in *T*_pull_final_ is shown alongside. **e**. Behavioral metrics of individuals in cluster A (*n* = 7) and B (*n* = 18) identified in (**d**) are shown. These plots are shown similarly in (**c**), but the colors indicate the respective cluster membership (magenta, cluster A; cyan, cluster B).

In providing the calcium activity from different cortical areas, Allen CCF was registered to the field of view by alignment of skull landmarks (MesoNet) ^36^ (Fig. 2e, Fig. S2a). We validated the accuracy of this method by comparing the MesoNet-based skull-landmark annotations with the manual annotations, based on three representative sessions from each of the 25 animals (Fig. 4a,b). In addition, the data from sensory mapping sessions further validated the accuracy of our mapping (Fig. 4c,d). In these sessions, we applied visual, auditory, and tactile sensory stimulation ^38–42^ to anesthetized mice (Fig. 4c). We obtained multimodal activity maps aligned onto the Allen CCF cortical map (Fig. 4d). Visual stimuli to the left eye of mice elicited prominent activity in the right occipital cortex ^39,40,43^ corresponding to VISp (primary visual area) in the atlas. Auditory stimuli evoked spreading activity, including a multimodal sensory area in the parietal cortex ^44^. Tactile stimuli to the right whisker pad elicited a robust response in the left SSp-bfd, a primary sensory cortex for processing whisker-related sensory information with marked hemispheric laterality ^45,46^. These specific activations of the cortical area validated the accurate mapping to the Allen CCF and also confirmed that the cortical calcium signals reflected neural activity. We also confirmed that the recorded animals did not show signs of aberrant short-duration, large-amplitude calcium activity, which has been reported by others ^28,47^ (Fig. S2b).

**Figure 4.**
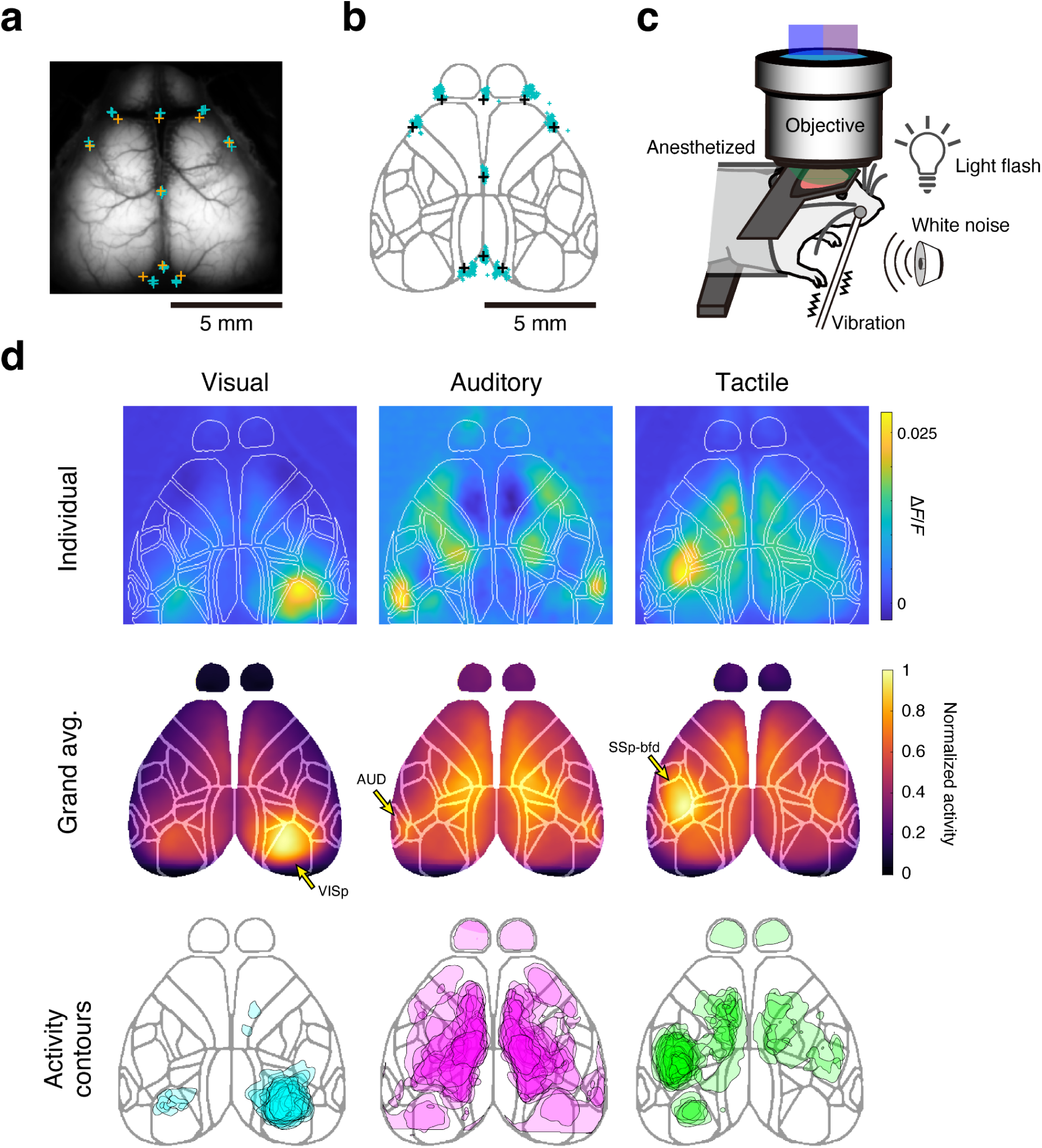
Validation of alignment to the Allen CCF reference atlas. **a,b**, Validation of MesoNet-based skull-landmark inference. In **a**, aligned skull landmarks from Allen CCF (orange) are plotted on the animal-average image of a representative animal, and are compared with the landmarks being manually annotated on three representative sessions from the animal (cyan). In **b**, manually annotated skull landmarks are warped and plotted on the Allen CCF atlas (cyan, 75 sessions from 25 animals), and compared with the reference landmarks (black). **c**, Schematic illustration of sensory mapping. Visual, auditory, and tactile stimulations were applied in the same task box as task training sessions (see Methods for details). **d**, Multi-modality sensory maps obtained from a representative mouse are shown (top panels). Color code shows normalized trial-averaged cortical responses (2 s duration from the stimulus onset) evoked by the three different sensory stimuli. Animal-averaged sensory maps of each modality (middle panels; *n* = 25 mice). Contours defined by more than 80-percentile value in the normalized map are displayed (Bottom panels). In each animal and each sensory stimulus, the evoked responses were normalized in the range from 0 to 1. Contours from 25 animals are overlaid. For this analysis, the data from the first session of sensory-mapping experiment in each animal are used.

In addition to calcium imaging data and task event timepoints, our dataset includes high-speed videography of the mice during every session. We took advantage of a widely used machine learning package for keypoint estimation, DeepLabCut ^12^, to estimate the position of the body part in each frame of videos from three directions, capturing the upper body, the right eye, and the right face (Fig. 5a–c). Comparison of DeepLabCut model-based keypoint annotations with manual annotations showed minor errors, validating the quality of our models (Fig. 5a-c). Moreover, the lever tip position estimated from the videography showed a markedly high correlation (0.98 ± 0.02; mean ± s.d., *n* = 356 sessions from 25 animals) with the encoder-based lever position (Fig. 5e,f). The position of the right forepaw used for pulling the lever should be correlated with the lever position, while the left forepaw should not. We found this to be exactly the case, based on the 357 task-recording sessions of our dataset that successfully included upper-body videos (Fig. 5d-f; right forepaw, 0.49 ± 0.23; left forepaw, 0.05 ± 0.19; mean±s.d., *n* = 356 sessions from 25 animals, also see Table 1; note that one out of the 357 available sessions did not contain any successful trials, and was omitted from the analysis).

**Figure 5.**
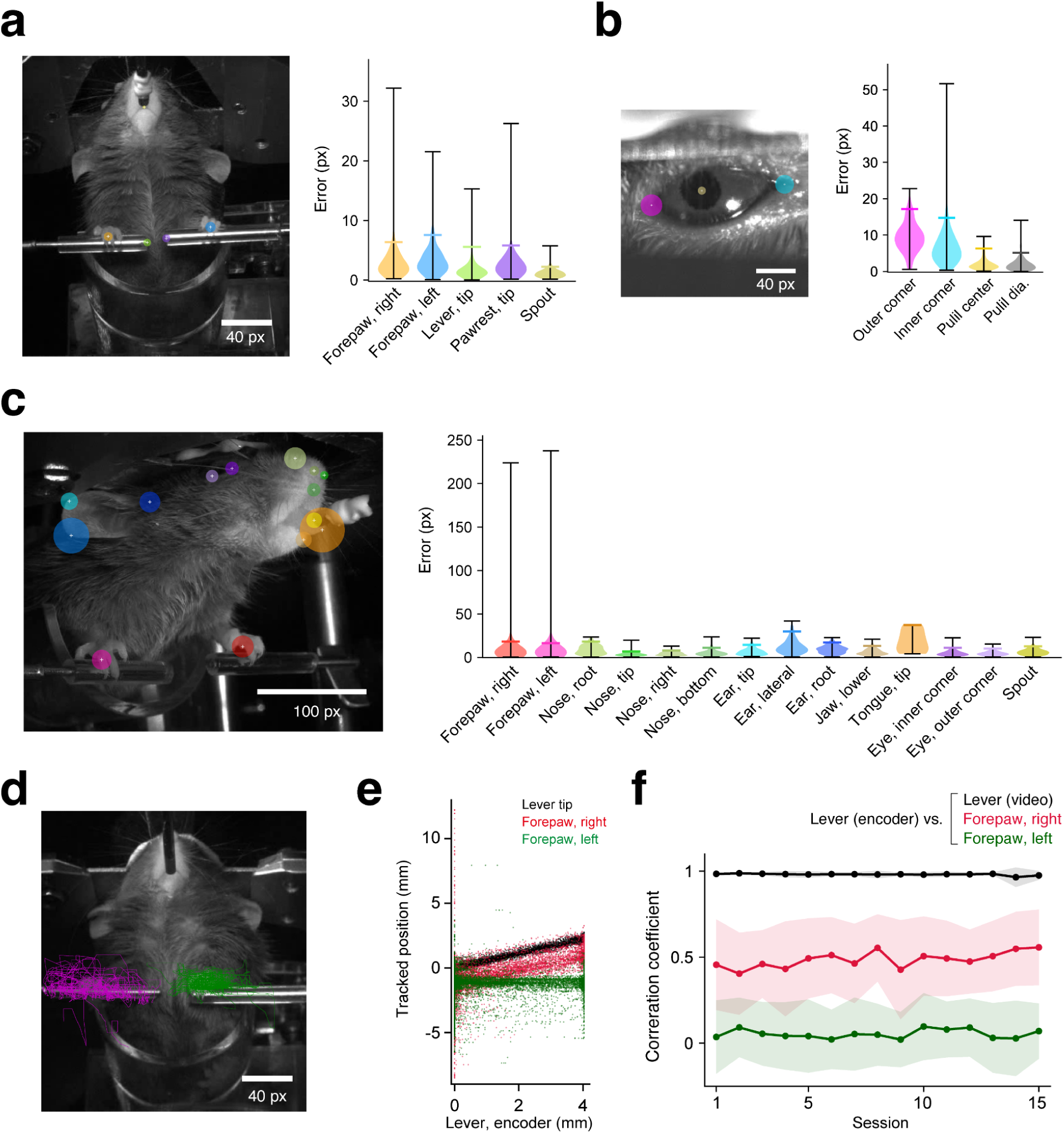
Validations of video-tracked keypoints. **a–c**, DeepLabCut-inferred and manually-annotated keypoints were compared using validation video frames (*n* = 192 frames from 25 animals for the body and the face cameras, and *n* = 190 frames from 25 animals for the eye camera, up to 8 frames per animal; see also Table 3). On the left panels, the 95 percentiles of the error distributions are indicated with colored circles overlaid on the manual annotations (white plus signs) in a representative frame of the body camera (**a**), the right eye (**b**), and the right face (**c**). The corresponding violin plots on the right panels are colored accordingly. Black whiskers indicate the minimum and the maximum errors, and colored horizontal lines represent the 95 percentiles. **d,e,** Keypoint positions identified as the lever tip, the right forepaws, and the left forepaws were compared with the lever position recorded by the rotary encoder, during all successful trials of a representative session. Keypoints identified in all trials are overlaid as lines (green; left, magenta; right) on a representative frame in (**d**), and their *y*-axis positions are plotted against the encoder-based lever position in (**e**) (black; lever tip, red; right forepaw, green; left forepaw). Each dot corresponds to a single frame of the videography. **f,** The Pearson’s correlation coefficient (*R*) between the keypoint positions and the encoder-based lever positions during successful trials was calculated across all the available task training sessions of all the animals (*n* = 25 mice, 356 sessions in total) and is displayed. The thick lines and shaded areas represent the mean±s.d. across the animals.

Finally, we analyzed cortical calcium activity and video-tracked body-part movements in conjunction with sensor-based behavioral measurements for an example task training session to validate the integrity of our data (Fig. 6). In this particular session, the robust synchrony among video-tracked movements, cortical activity, and task events is apparent even in raw traces (Fig. 6a), and becomes more distinct when these signals are aligned to the lever-pull onset (Fig. 6b) ^17,19,48–51^. While some body parts, such as forepaws and nose, began moving before the lever-pull onset, others, including jaw and pupil diameter, moved afterward at least in this session. Aligned traces of many cortical areas show onsets similar to lever-pull movements, and motor and somatosensory areas showed the activity relatively correlated with lever-pull movements. We found slight differences between them. The secondary motor cortex (MOs) quickly recovered to the baseline level. In contrast, the primary motor (MOp) and somatosensory cortex (SSpul and SSpn) showed slightly higher activity than the baseline level after the reward delivery. Other sensory-related areas (OB, olfactory bulb; RSPagl, retrosplenial area, lateral agranular part; AUDs, secondary auditory areas; VISp, primary visual area) showed activity traces that shared time courses closely matching the pupil dilation. The data also revealed a decrease in activity variability after the lever-pull onset, exemplifying how stereotypic movements like lever-pull constrain cortical activity. Furthermore, since our cortical activity data inherently contains spatial information and has been preprocessed to a ready-to-use format, the spatiotemporal dynamics of cortical activity are readily accessible (Fig. 6c).

**Figure 6.**
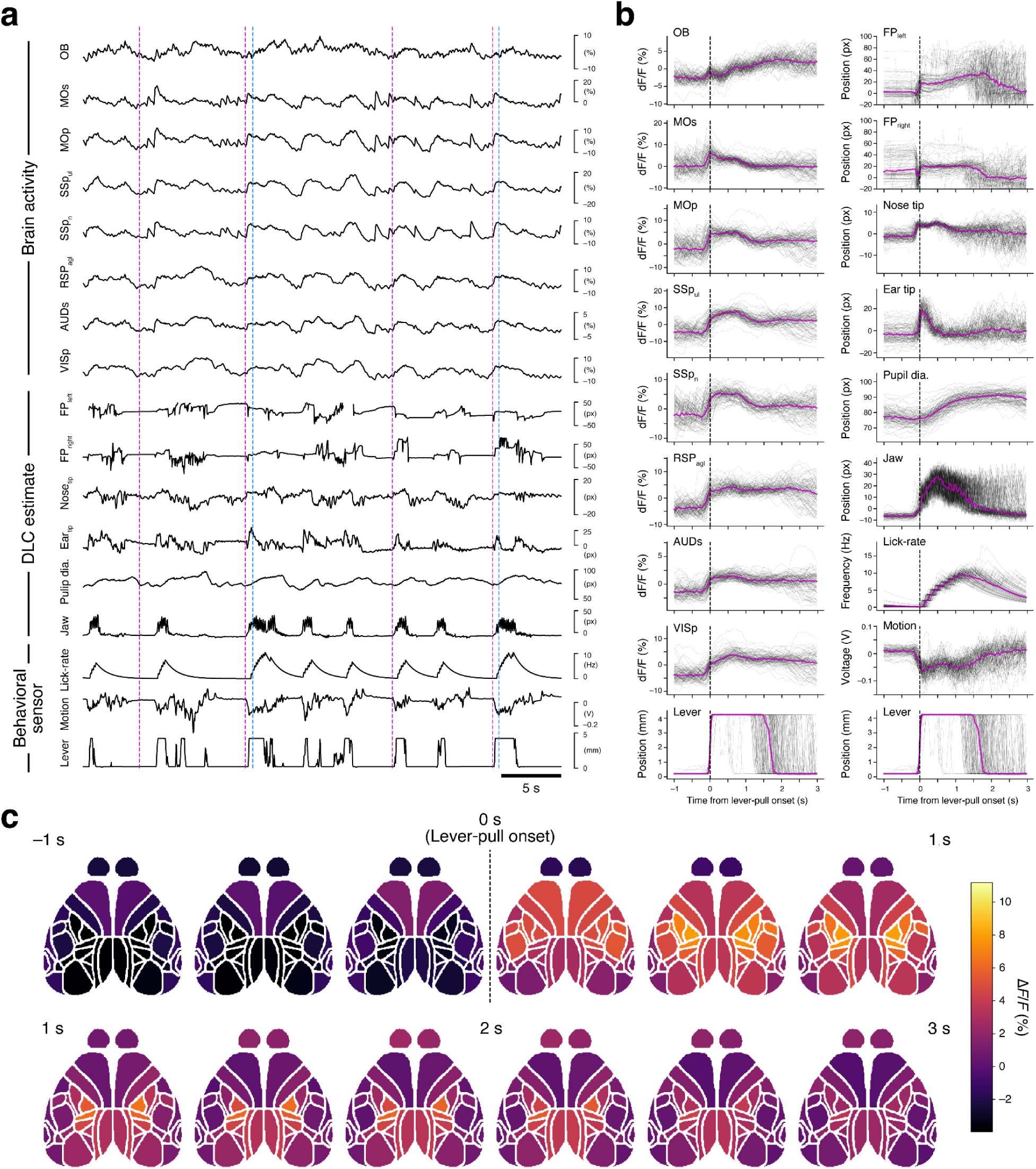
Validations of task-related behavior and activity in a representative session. **a**, Raw trajectories of calcium dynamics from various cortical regions and video-tracked body-part movements with sensor-based behavioral records (lever movement, gross body motion, and lick rate), referenced to task events (magenta, sound cue onset; blue, reward delivery). The disconnected part in the trace of the left forepaw is caused by the value deficit with the deviation from the 5 and 95 percentiles of DLC estimation. **b**, Lever-pull aligned traces of calcium dynamics (left column) as well as video-tracked body-part movements and sensor-based behavioral records (right column). Trial-averaged (magenta) and single-trial (black) traces from a representative session are shown. The bottom-most traces of both columns show the same lever traces as references. **c**, Trial-averaged calcium activity was mapped onto the Allen CCF and shown in chronological order from the top-left to the bottom-right. Calcium activity was binned into intervals of 10 frames (333 ms) and each bin was averaged. The labels (−1 s, 1 s, 2 s, 3 s) show time relative to lever-pull onset. The data are from the same session shown in (**a**) and (**b**).

## Code availability

All workflows presented in this paper were implemented in MATLAB and Python, and the source code is publicly available (MIT License) at: https://github.com/BraiDyn-BC/bdbc-data-pipeline.

## Acknowledgements

We thank M. Nishiyama for animal care, and S.-I. Terada for help setting up optical equipment in the one-photon calcium imaging device and the technical advice. This work was supported by JSPS KAKENHI under Grant Number JP22H05163 to K.N.; JP22H05160, JP22H05154 to M.M.; JP22H05162, JP23K20309 to Y.R.T., AMED under Grant Number JP19dm0207088 and JP24wm0625404 to K.N.; JP24wm0625001 to M.M; JP24wm0625113 to Y.R.T., JST under Grant Number JPMJMS2294 to Y.R.T., and the Cooperative Study Program of Exploratory Research Center on Life and Living Systems (ExCELLS; program no. 19-102 to K.N.).

## Author Contributions

M.K. was responsible for data curation, investigation, methodology, software and visualization. K.S. was responsible for data curation, formal analysis, software, validation and visualization. R.H. was responsible for investigation and resources. R.A. are responsible for investigation, resources and validation. S.S. carried out NWB formatting within the data pipeline. Y.R.T. contributed to conceptualization, funding acquisition, supervision and manuscript writing. M.M. contributed to conceptualization, funding acquisition, project administration, and manuscript writing. K.N. contributed to conceptualization, funding acquisition, project administration, resources, supervision, visualization and manuscript writing. The contribution diagram is shown in Fig. S4.

## Corresponding authors

Correspondence to Masashi Kondo and Ken Nakae.

## Author Information

These authors contributed equally: Masashi Kondo, Keisuke Sehara, Rie Harukuni and Ryo Aoki.

---

**Figure S1.**
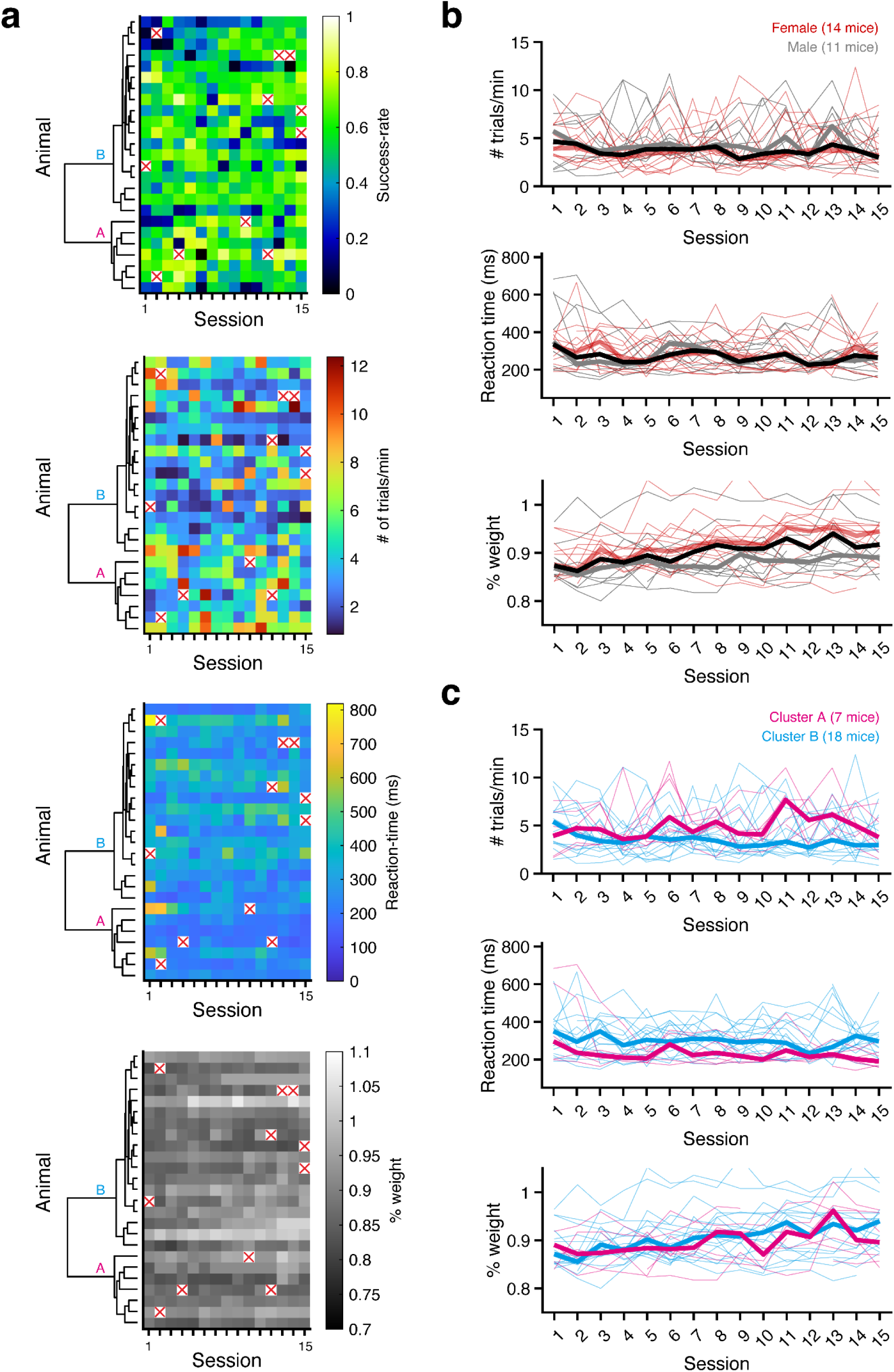
Behavioral performance in task training sessions. **a**, From top to bottom, four different behavioral parameters obtained are color-coded and shown. **b**, Trial rate (The trial number per min; top panel), reaction time (in all lever-pulled trials; middle panel), body weight (ratio to the weight before starting water restriction; bottom panel) across sessions are shown. Thin lines show performances of individual animals (red, female; gray, male), whereas thick lines are the median across the population (red, female; gray, male; black, all). **c**, Behavioral parameters of individuals in cluster A (*n* = 7) and B (*n* = 18) identified in Fig. 3d are shown. These plots are shown similarly in (**b**), but the colors indicate the respective cluster membership (magenta, cluster A; cyan, cluster B).

**Figure S2.**
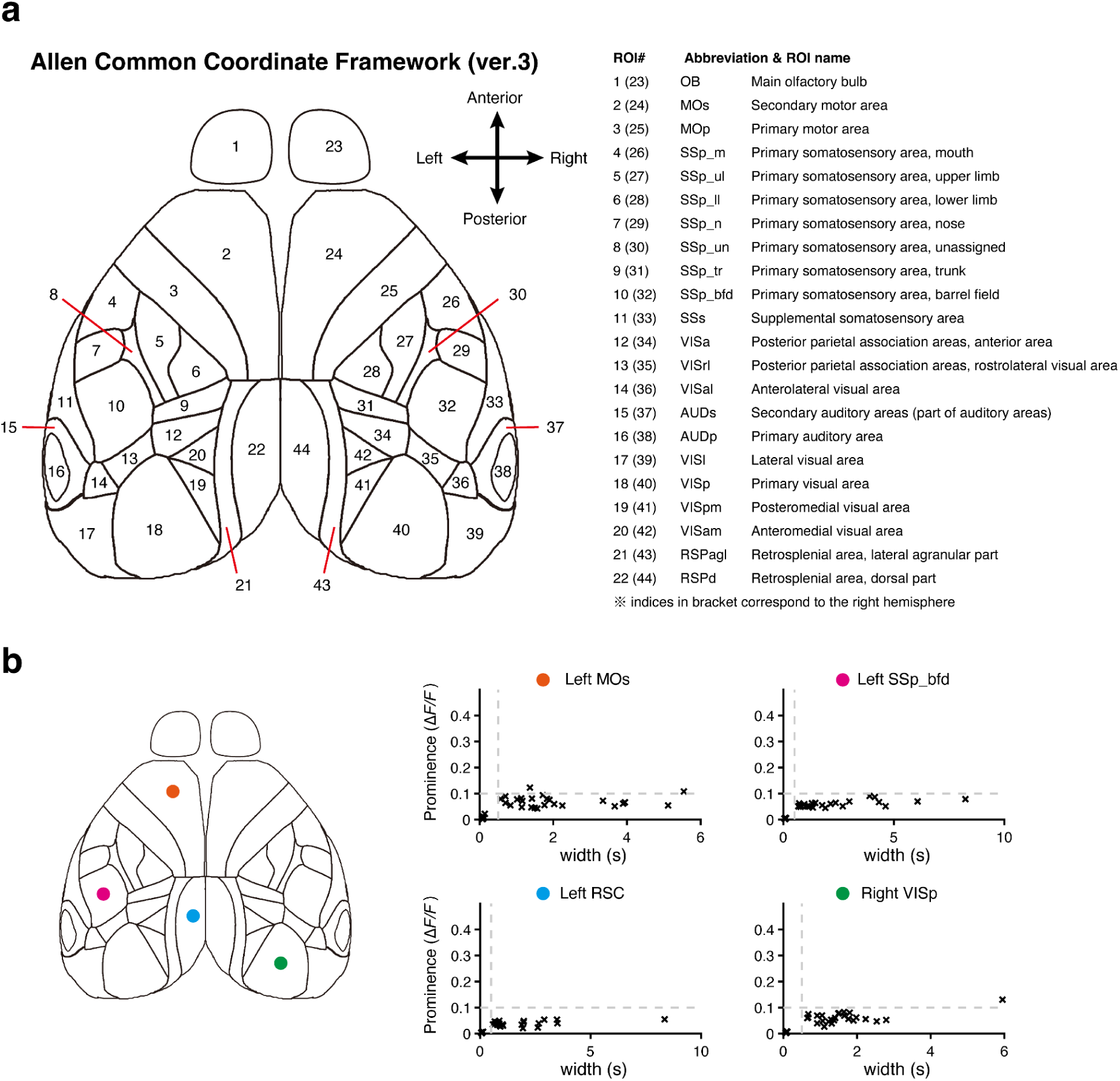
**a,** Coordinates and names of segmented brain regions used for the dataset. It originated from the dorsal view of the Allen Common Coordinate Framework (version 3). **b,** The validation of aberrant activity in the transgenic mouse was used in this study. Prominences and widths of calcium activities were detected in each first resting-state recording of each animal. The left side of frontal, barrel, retrosplenial, and right side of visual regions were selected for the analysis (left). These were clustered with K-means clustering (k = 2) in each animal and region, and then the centers of mass of two clusters were obtained. High prominence (> 0.1 in Δ*F*/*F*) and narrow width (< 0.5 s) of the transients were not detected in each animal (right).

**Figure S3.**
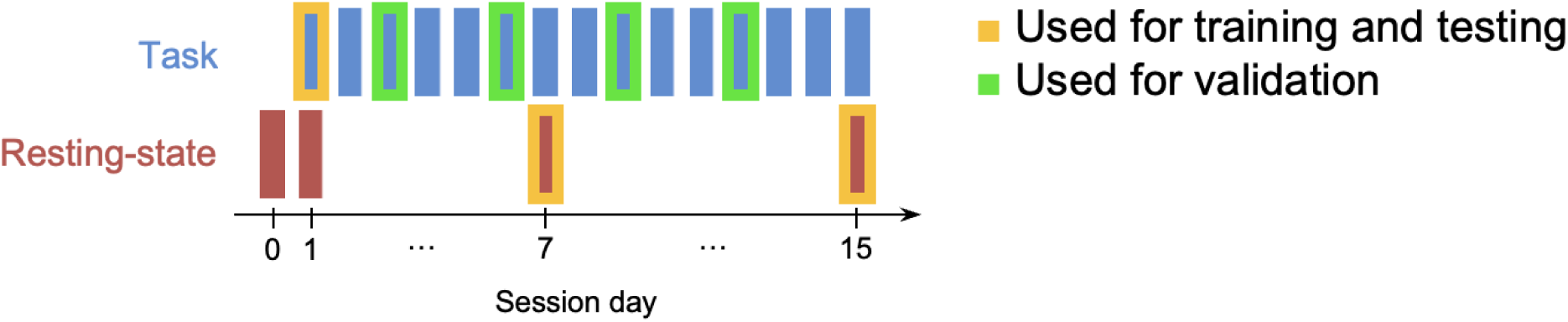
Schematic of frame extraction strategy for training and validating DeepLabCut models. Task training session 1 and resting-state recording sessions 7 and 15 (including the day-8 resting-state session from one animal) were used to extract video frames for training and testing. Validation frames, for comparison with manual annotation, were extracted from task training sessions 3, 6, 9, and 12.

**Figure S4.**
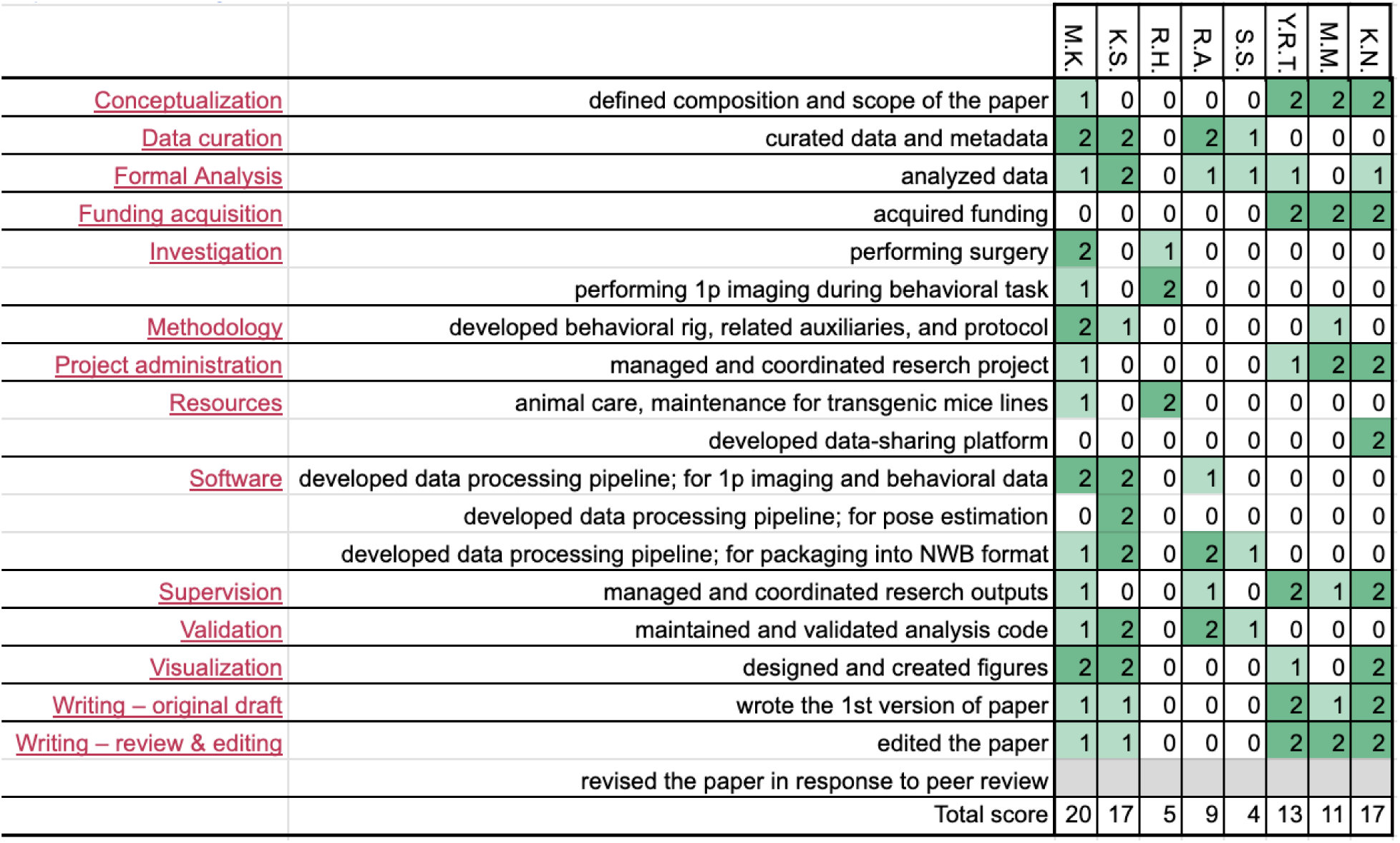
Contribution diagram. The following diagram illustrates the contributions of each author, based on the CRediT taxonomy (Brand et al., 2015). For each type of contribution there are three levels indicated by color in the diagram: ’support’ (light) and ‘lead’ (dark).

